# Inter- and intrapopulational heterogeneity of characteristic markers in adult human neural crest-derived stem cells

**DOI:** 10.1101/2021.06.25.449886

**Authors:** Beatrice A. Windmöller, Anna L. Höving, Johannes F.W. Greiner

**Affiliations:** Department of Cell Biology, University of Bielefeld, Bielefeld, Germany; Forschungsverbund BioMedizin Bielefeld FBMB e.V., Bielefeld, Germany

**Keywords:** Heterogeneity, adult stem cells, neural crest-derived stem cells, inferior turbinate stem cells, stochastic heterogeneity, clonal heterogeneity

## Abstract

Adult human neural crest-derived stem cells (NCSCs) are found in a variety of adult tissues and show an extraordinary broad developmental potential. NCSCs are therefore promising candidates for applications in regenerative medicine, although increasing evidence suggest a remaining niche-dependent variability between different NCSC-populations regarding their behavior and expression signatures. In the present study, we extended the view on heterogeneity of NCSCs by identifying heterogeneous expression levels and protein amounts of characteristic markers even between NCSCs from the same niche of origin. In particular, populations of neural crest-derived inferior turbinate stem cells (ITSCs) isolated from different individuals showed significant variations in characteristic NCSC marker proteins Nestin, S100 and Slug in a donor-dependent manner. Notably, increased nuclear protein amounts of Slug were accompanied by a significantly elevated level of nuclear NF-κB-p65 protein, suggesting an NF-κB-dependent regulation of NCSC-makers. In addition to this interpopulational genetic heterogeneity of ITSC-populations from different donors, single ITSCs also revealed a strong heterogeneity regarding the protein amounts of Nestin, S100, Slug and NF-κB-p65 even within the same clonal culture. Our present findings therefor strongly suggest ITSC-heterogeneity to be at least partly based on an interpopulational genetic heterogeneity dependent on the donor accompanied by a stochastic intrapopulational heterogeneity between single cells. We propose this stochastic intrapopulational heterogeneity to occur in addition to the already described genetic variability between clonal NCSC-cultures and the niche-dependent plasticity of NCSCs. Our observations offer a novel perspective on NCSC-heterogeneity, which may build the basis to understand heterogeneous NCSC-behavior and improve their clinical applicability.

## Introduction

Stem cells with an embryonic origin from the neural crest crucially participate in embryonic development, but are also commonly known to remain within the organism until adulthood, where they reside as quiescent adult stem cells. During development, the neural crest (NC) arises between the future ectoderm and the neural tube following the process of neurulation ^1^. After undergoing epithelial to mesenchymal transition (EMT), neural crest cells (NCCs) migrate in range of developing tissues and contribute to development by guiding patterning and by differentiation. Amongst others, NCCs give rise to most parts of the craniofacial skeleton, enteric neurons of the foregut and stomach or sympathetic ganglia and melanocytes in the trunk (reviewed in ^2–4^). After participating to embryonic development, pools of NCCs remain as adult neural crest-derived stem cells (NCSCs) in various tissues including the skin ^5^, hair follicles ^6^, carotid body^7^, heart ^8,9^, palatum ^10^ or nasal cavity ^11–13^ (reviewed in ^2,3^). Adult NCSCs share a broad differentiation potential particularly into mesodermal and ectodermal derivates as well as the expression of characteristic markers. These marker proteins include the intermediate filament Nestin and the calcium binding protein S100 as well as NC-related transcription factors (TF) like Twist or Slug, which belong to group of TF enabling EMT and therefore migration of embryonic NCCs (reviewed in ^2,3^). Their developmental potential and accessibility makes NCSCs promising candidates for regenerative medicine ^14,15^ and pharmacological research ^16,17^. Despite similarities in differentiation capacity and marker expression, increasing evidences show a remaining heterogeneity between different NCSC-populations (reviewed in ^2^). In this regard, we very recently observed an interpopulational heterogeneity between craniofacial and cardiac NCSCs. Although we demonstrated high similarities between the global transcriptomes of both NCSC-populations, gene expression patterns and differentiation potentials differed with particular regard to their tissue of origin^8^. Here, cardiac stem cells showed an exclusively high capacity for differentiation into cardiomyocytes, while craniofacial NCSCs more efficiently gave rise to neurons or osteoblast compared to their cardiac counterparts ^8^. These observations are in line with several studies reporting an association between the differentiation behaviors of NCSCs and their niche. For instance, NCSCs from the murine carotid body were demonstrated not only to undergo neurogenesis but even angiogenesis i*n vivo* ^7,18^. On the contrary, NC-derived dental pulp stem cells showed the capacity to form dentin and dental pulp tissue in addition to their neuronal and osteogenic differentiation potential ^19^, suggesting a niche-dependent plasticity of adult NCSCs.

In the present study, we extended this perspective on NCSC-heterogeneity by reporting a highly heterogeneous marker expression even between and within NCSC-populations from the same niche of origin. We particularly took advantage of an adult craniofacial NCSC-population isolated from the inferior turbinate of the nasal cavity (inferior turbinate stem cells, ITSCs)^13^, which we previously utilized for the interpopulational comparison with cardiac NCSCs^8^. Like other NCSC-populations, ITSCs show a broad differentiation potential including the capacity to give rise to glutamatergic and dopaminergic neurons ^15,20^ or osteoblasts ^21^ as well as the expression of common NCSC-markers^13,22^. However, our present data reveal a broad heterogeneity between ITSCs regarding the expression and protein amounts of such NCSC-markers, particularly Nestin, S100, Twist and Slug. On an interpopulational level, the protein amounts of Nestin, S100 and Slug significantly differed between ITSC-populations from distinct donors. Notably, significantly elevated nuclear protein levels of Slug in one female donor were accompanied by a significantly increased amount of nuclear factor kappa-light-chain-enhancer of activated B cells (NF-κB) protein in ITSCs. NF-κB is a key regulator of various cellular processes including cell growth, inflammation, memory or immunity^23–25^ and was recently described to even control fate choices of ITSCs^26^. As for Nestin, S100 and Slug, the amount of nuclear NF-κB-p65 protein within a population of ITSCs was found to be highly dependent on the donor. In addition to this interpopulational heterogeneity, we observed a remarkable intrapopulational heterogeneity of Nestin, S100, Slug and NF-κB-p65 protein levels between single ITSCs within a respective population and even within clonally grown cultures.

## Materials and Methods

### Cell culture

ITSCs were isolated from inferior turbinate tissue obtained during routine nasal surgery according to Hauser and coworkers^13^ after an informed consent according to local and international guidelines. All experimental procedures were ethically approved by the ethics board of the medical faculty of the University of Münster (No. 2012–015-f-S). Cells were cultured in Dulbecco’s modified Eagle’s medium/Ham F-12 (DMEM-F12) (Sigma Aldrich, St. Louis, MO, USA) supplemented with 40 ng/ml basic fibroblast growth factor-2 (FGF2) (Miltenyi Biotec, Bergisch Gladbach, Germany), 20 ng/ml epidermal growth factor (EGF) (Miltenyi Biotec), 1x B27-Supplement (Thermo Fisher Scientific, Waltham, MA, USA), 200 mM L-glutamine (Sigma Aldrich), 10 mg/ml Penicillin/Streptomycin (Sigma Aldrich) and 0,25 mg/ml Amphotericin B (Sigma Aldrich) with additional 10% human blood plasma (Institute for Laboratory- and Transfusion-Medicine, Heart- and Diabetes-Centre NRW, Bad Oeynhausen, Germany) at 37°C, 5% O_2_ and 5% CO_2_ in a humidified incubator (Binder, Germany) as previously described ^22^. Passaging was performed using Collagenase I (Sigma-Aldrich) as previously described^13,22^.

### Transfection of cultivated ITSCs

ITSCs cultivated as described above were enzymatically detached using collagenase I and harvested at 300 x g for 10 min. Dissociated cells were transfected with 1 μg pmax GFP vector (Amaxa Biosystems, Switzerland) and Amaxa rat NSC-Nucleofector Kit (Amaxa Biosystems) and the Nucleofector II device (Amaxa Biosystems) according to the manufacturer’s information. 5 ml DMEM F-12 (Sigma Aldrich) were added before the sample was centrifuged for 10 min at 300 x g. After discarding the supernatant, cells were cultured as described above. Imaging of transfected cells was performed using a Axio Oberver.D1-Microscope (Carl Zeiss AG, Oberkochen, Germany).

### Flow cytometry and FACS

Flow cytometric analysis and fluorescence activated cell sorting (FACS) of GFP-transfected ITSCs was performed using CyFlow Space (Partec, Münster, Germany). 10,000 GFP^+^ cells were sorted and collected in 15 ml reaction tubes (Sarstedt, Nümbrecht, Germany) and subsequently stored on ice for further experimental procedures. Data were processed using FlowJo software (Tree Star, OR, USA).

### SMARTseq2

After the isolation of single cells, SMARTseq2 was performed according to Picelli and colleagues ^27^. Briefly, cells were transferred in a total volume of 0.5 μl into 2 μl of cell lysis buffer consisting of 0.2 % (v/v) Triton X-100 (Sigma Aldrich) and 2 U/μl RNase Inhibitor (Invitrogen, Carlsbad, CA, USA) with additionally 1 μl oligo-dT primers and 1 μl of dNTP mix (NEB Biolabs, Ipswich, MA, USA). Further, hybridization of the oligo-dT primers to the poly (A) tail of the mRNA, first-strand-synthesis with template switching and PCR-preamplification were performed as described by Picelli and coworkers ^27^.

### RT-PCR

PCR was performed using GoTaq DNA polymerase (Promega, Fitchburg, WI, USA) according to the manufacturer’s guidelines with the following primers for eEF2 (AGGTCGGTTCTACGCCTTTG, TTCCCACAAGGCACATCCTC), Vimentin (GTGGACCAGCTAACCAACGACAAA, AGGTCAGGCTTGGAAACATCCACA), GAPDH (CATGAGAAGTATGACAACAGCCT, AGTCCTTCCACGATACCAAAGT), S100 (GGGAGACAAGCACAAGCTGAAGA, TCAAAGAACTCGTGGCAGGCAGTA), Snail (CCCAATCGGAAGCCTAACTA, GGACAGAGTCCCAGATGAGC) and Twist (GTCCGCAGTCTTACGAGGAG, CCAGCTTGAGGGTCTGAATC)

### Immunocytochemistry

For Immunocytochemistry, ITSCs cultivated as described above were seeded on top of cover glasses in and cultivated for two days followed by fixation with phosphate-buffered 4 % paraformaldehyde (lab-made) for 15 min at room temperature (RT). Permeabilization and blocking was subsequently done using 0,002 % Triton X-100 (Sigma-Aldrich, Germany) and 5 % goat serum (Dianaova, Germany) or 1 % BSA (in PBS) diluted for 30 min at RT. Primary antibodies against Nestin (Mouse, 10C2, 1:300, Merk KGaA, Germany), S100 (Rabbit, Z0311, 1:400, Dako Cytomation, Germany), Slug (Rabbit, C19G7, 1:100, Cell Signaling, Germany) or NF-κB-p65 (Mouse, 200301065, 1:5000, Rockland, PA USA) were incubated for 1 h at RT. Secondary fluorochrome-conjugated antibodies (Alexa 555 anti-mouse, Alexa 555 anti-rabbit, Alexa 488 antimouse, 1:300, Life Technologies, Germany) were incubated for 1 h at RT followed by nuclear counterstaining with DAPI (1 μg/ml in 1 x PBS; Sigma-Aldrich, Germany) for 10 min at RT. Fluorescence imaging was done with a confocal laser scanning microscope (LSM 780; Carl Zeiss, Jena, Germany). Five randomly placed pictures were taken per donor and analyzed using ImageJ/Fiji ^28^.

## Results

### 1. Transcriptional profiling of single adult human inferior turbinate stem cells reveals strong differences in the expression of NC-specific genes

To initially assess the potential heterogeneity of inferior turbinate stem cells isolated from the human nasal cavity (Figure 1A) on transcriptional level, we successfully applied SMARTseq2^27^ on single ITSCs. To facilitate a fluorescence-based sorting procedure, ITSCs were transfected with GFP (Figure 1B). Analysis of GFP^+^-cells during flow cytometric sorting revealed a transfection efficiency of 23,2 % (Figure 1C,D). SMARTseq2 was performed with samples comprising ten cells and three samples consisting of one cell. A following RT-PCR analysis revealed the presence of transcripts for GAPDH, eEF2 and Vimentin in the ten-cells-sample as well as in all three single cell approaches (Figure 1 E). Further transcriptional profiling revealed the heterogenous expression of the NCSC-markers S100, Snail and Twist, the latter being key players in EMT and highly important in the determination of NCSC-fate (reviewed in ^2^). In particular, S100 gene expression was detectable in the 10-cell-sample and in two out of three single-cell-samples. Interestingly, Snail transcripts could not be observed within the 10-cell-sample but in two of the single ITSCs. Notably, Twist mRNA could be successfully amplified in the sample originating from ten ITSCs as well as in all three single-cell-samples with varying expression level (Figure 1E). These data indicate a general transcriptional heterogeneity between single ITSCs from the same population.

**Figure 1:**
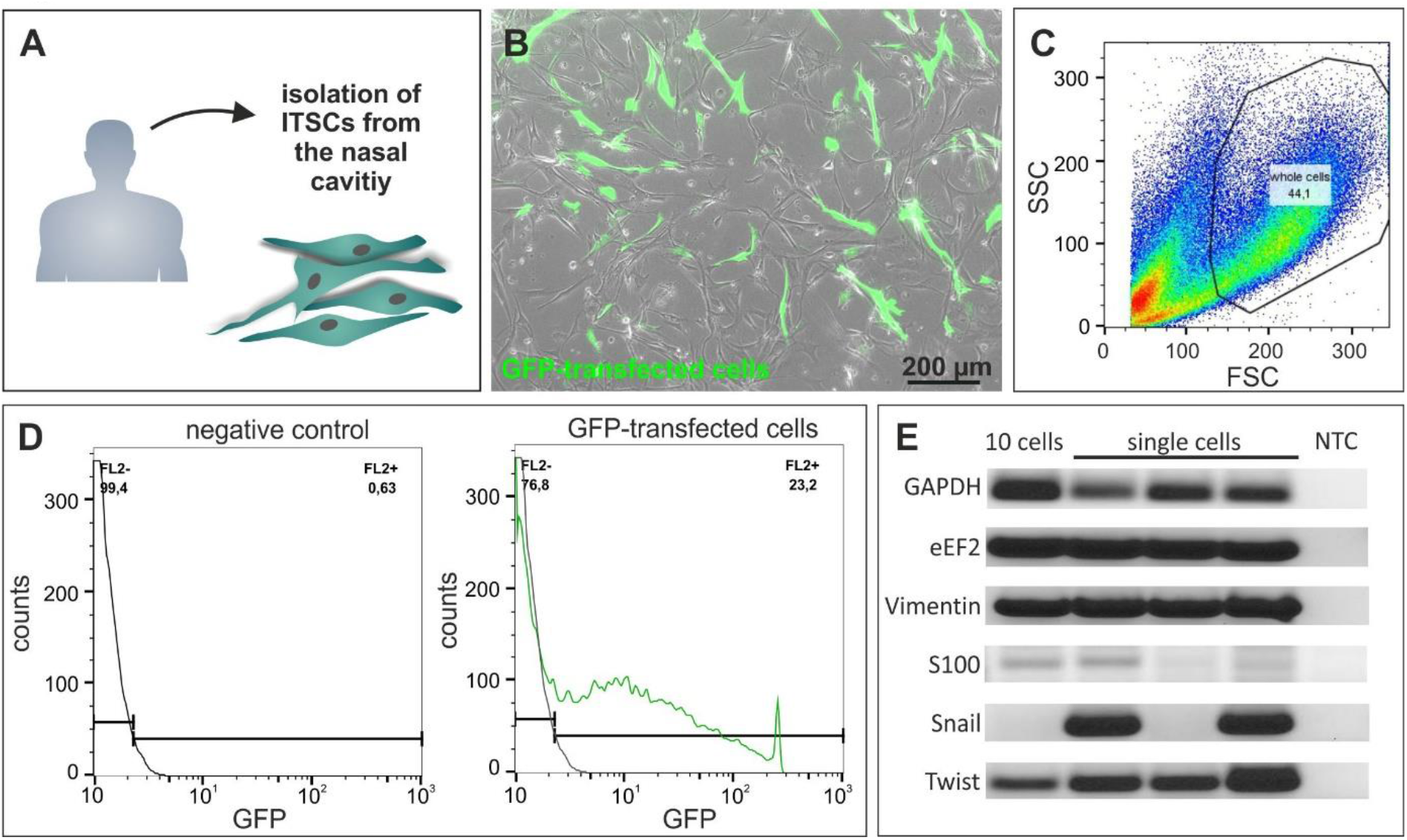
SMARTseq2 reveals strong heterogeneity of single ITSCs in their NCSC-marker expression. A: ITSCs were derived from the inferior turbinate of the human nasal cavity. B: GFP-transfected ITCSs 24 h after transfection. C: Flow cytometric analyses of ITSCs. SSC and FSC measure size and granularity of the cells. Whole cells (44.1 %) are marked for further gating. D: Sorting of GFP^+^ ITSCs. FL2 measures emitted fluorescence signal (GFP). Untransfected cells serve as negative control. E: RT-PCR after SMARTseq2 for the housekeeping genes GAPDH, eEF2 and Vimentin as well as for the NCSC-markers S100, Snail and Twist.

### 2. ITSCs show highly heterogeneous protein levels of Nestin and S100 in a donor-dependent manner

To investigate if the heterogeneous expression of neural crest and stem cell markers in single ITSCs observed on mRNA level is also present on protein level, immunocytochemistry against Nestin and S100 was performed using ITSCs isolated from three male and three female donors (Figure 2A). Here, the intermediate filament and stem cell marker Nestin displayed a strong heterogeneity on protein level within the respective ITSC-populations (Figure 2B-C). Single ITSCs differed between low (Figure 2B-C, arrowheads) and high (Figure 2B-C, arrows) amounts of Nestin protein. Notably, we nevertheless observed significant differences in the amount of Nestin protein between ITSC-populations from distinct donors in a sex-independent manner (Figure 2D). A quantification of the fluorescence intensities also revealed intrapopulational variations in the amount of the neural crest-related protein S100, which was further shown to be significantly varying between ITSCs-population in dependence to the donor (Figure 2 E-G).

**Figure 2:**
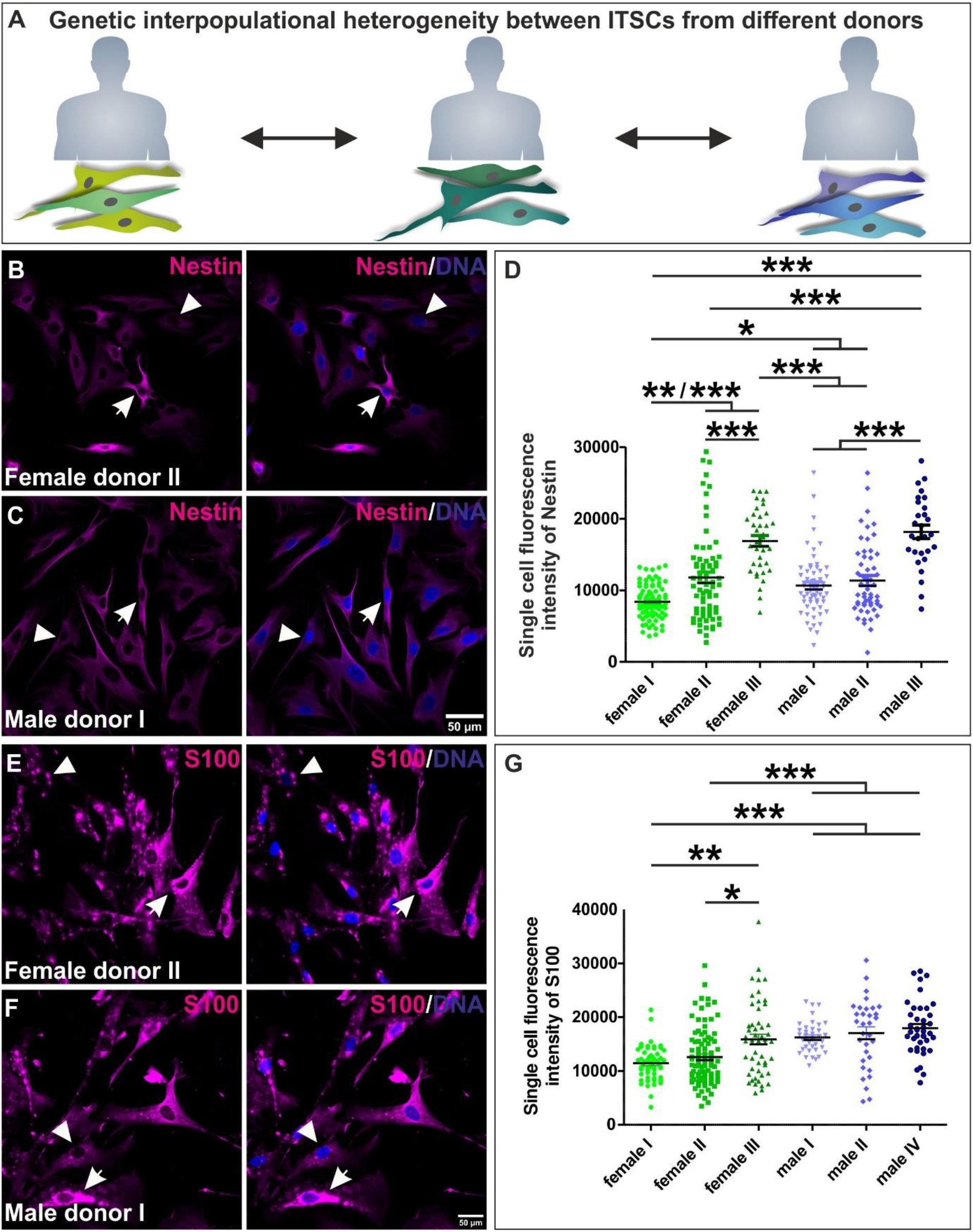
Immunocytochemistry confirmed heterogeneous expression of Nestin and S100 between ITSCs in dependence to the donor. A: Schematic view on interpopulational heterogeneity in stem cell marker expression between individual ITSC-populations in dependence to the donor. B-C: Representative images of immunocytochemical stained Nestin protein in female and male-derived ITSCs. D: Quantification of immunocytochemistry via measurement of the cytosolic fluorescence intensity of Nestin per single cell validated a significant heterogeneity dependent on the donor on protein level. E-F: Representative images of immunocytochemical stained S100 protein in female and male-derived ITSCs. G: Quantification of immunocytochemistry via measurement of the cytosolic fluorescence intensity of S100 per single cell affirmed the heterogeneity of S100 expression on protein level. Kruskal-Wallis test, Post test: Dunn’s Multiple Comparison Test, *p<0.05, *p<0.01, *** p < 0.001. was considered significant.

In conclusion, we observed a great donor-dependent heterogeneity in the amounts of Nestin and S100 protein between ITSC-populations, although protein amounts likewise strongly varied between single ITSCs.

### 3. Variations in nuclear localized Slug and NF-κB-p65 proteins are present between ITSCs on intrapopulational and interpopulational level with regard to the donor

Assessing potential variation of regulatory NC-related transcription factors like Slug in ITSCs, we observed a strong heterogeneity between single ITSCs within a distinct population regarding the amount of nuclear localized Slug protein (Figure 3A-B). Measurements of the nuclear fluorescence intensities further revealed a significant heterogeneity between the different ITSC-populations related to the donor but in a sex-independent manner (Figure 3C). With regard to the already described regulation of Slug by NF-κB^29^ and its very recently described role in differentiation of ITSCs^30^, we also determined NF-κB-p65 nuclear protein amounts in single ITSCs. Likewise to our observation regarding Slug protein, we observed a strong intra- and interpopulational heterogeneity of NF-κB-p65 protein amounts between ITSCs (Figure 3A-C). Interestingly, the ITSC-population isolated from female donor II showing a significantly increased nuclear protein amount of Slug also revealed a significantly elevated level of nuclear localized NF-κB-p65 (Figure 3C). These findings are in line with the recently described correlation between the activity of NF-κB-p65 and Slug^29^.

**Figure 3:**
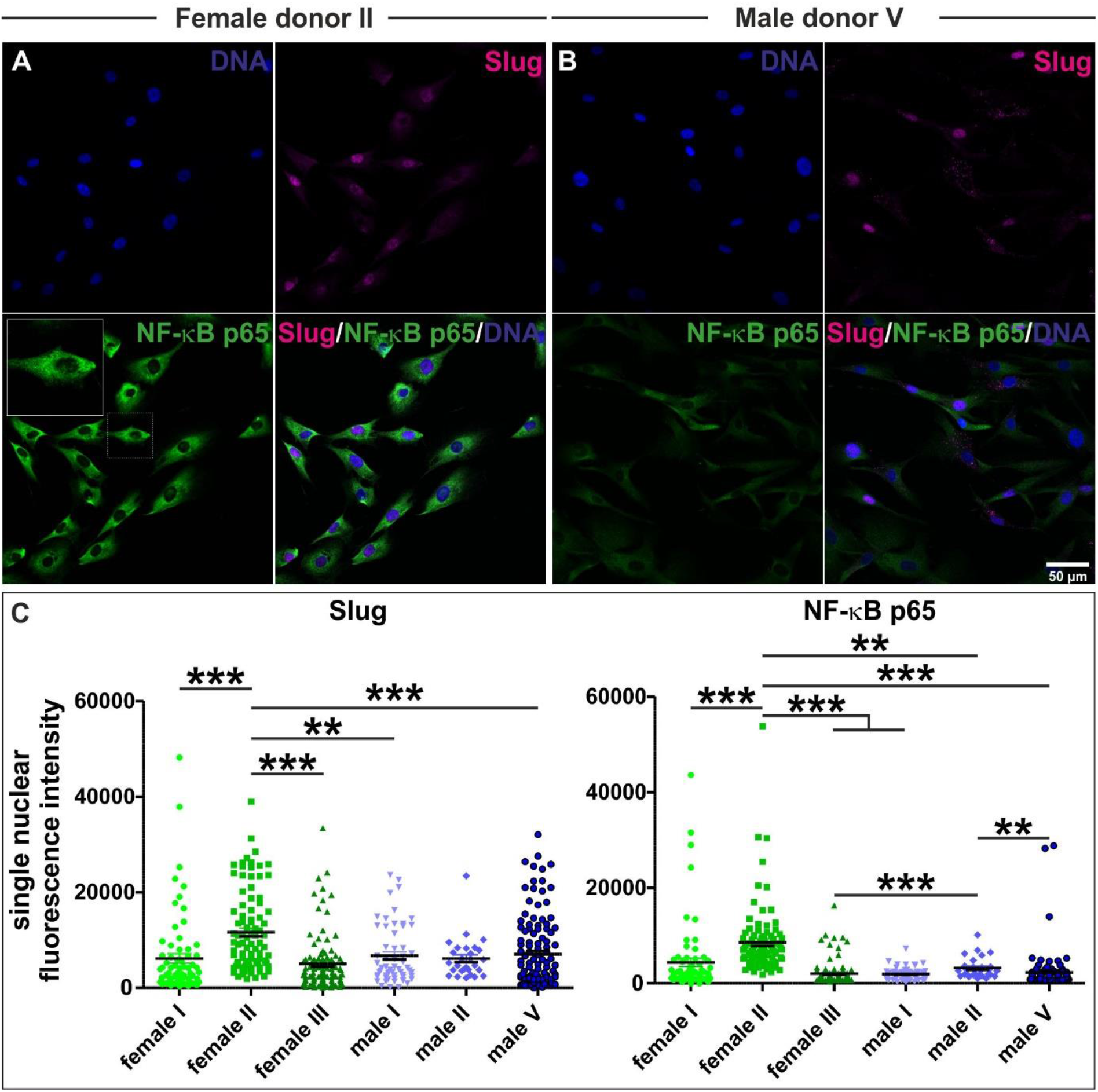
Intra- and interpopulational heterogeneity of NF-κB-p65 and the EMT transcription factor Slug. A-B: Representative images of immunocytochemical stainings showing heterogeneity of Slug and NF-κB-p65 protein between single ITSCs and ITSC-populations. C: Quantification of immunocytochemical stainings via measurement of the nuclear fluorescence intensity of Slug and NF-κB-p65 per single cell revealed an intrapopulation and interpopulational variability with regard to the donor. Kruskal-Wallis test, Post test: Dunn’s Multiple Comparison Test, *p<0.05, *p<0.01, *** p < 0.001. was considered significant.

### 4. Heterogeneous protein amounts of neural crest and stem cell markers are also observable between single ITSCs within clonally grown cultures

To assess the origins of the heterogeneous expression of neural crest and stem cell markers observed between single ITSCs, we determined the protein amounts of Nestin, S100, Slug and NF-κB-p65 in clonally grown ITSCs (Figure 4 A). Contrary to our assumption of a more homogenous distribution of protein amounts, quantification of the immunofluorescence intensities on single cell level revealed highly heterogeneous protein levels of the stem cell marker Nestin between ITSCs in the same clonal culture (Figure 4 B-C). Although the NCSC-marker S100 was expressed in high amounts within all analyzed ITSCs of the clonal culture, the protein amount differed between single cells (Figure 4 D-E). Moreover, nuclear fluorescence intensity of Slug and NF-κB-p65 showed highly variable degrees of nuclear translocation on single cell level between ITSCs within a clonal population (Figure 4 F-I). In summary, we provide evidence for a strong heterogeneity even between single ITSCs in the same clonal culture regarding the protein levels of neural crest and stem cell markers as well as of the subunit p65 of the NF-κB family.

**Figure 4:**
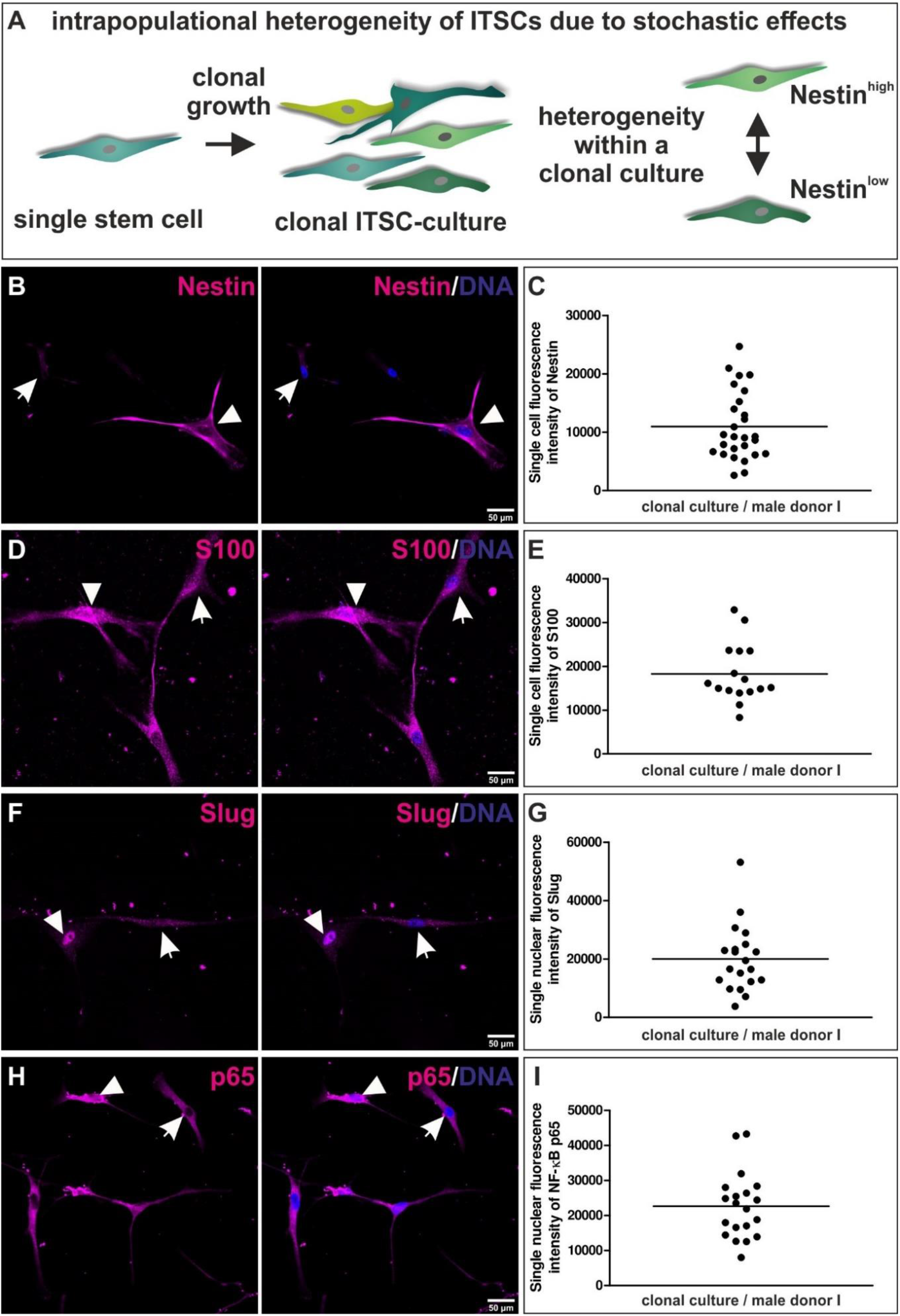
Stochastic single cell heterogeneity within clonally grown ITSC-cultures. A: Schematic view on heterogenous protein amounts between single ITSCs in the same clonal culture. B, D, F, H: Representative images of immunocytochemical stainings showing heterogeneity of Nestin, S100, Slug and NF-κB-p65 proteins between single ITSCs present in the same clonally grown culture. C, E, G, I: Quantification of immunocytochemical stainings via measurement of the single cell or single nuclear fluorescence intensity showing the stachostic variability in Nestin, S100, Slug and NF-κB-p65 protein amounts.

## Discussion

The present study reveals a new perspective on the heterogeneity of adult human neural crest-derived stem cells by demonstrating donor-dependent interpopulational variations in NCSC-marker genes, which are accompanied by an intrapopulational heterogeneity even within clonal cultures.

On the level of donor-to-donor variabilities, we found the protein amounts of Nestin, S100 and Slug to be significantly different between populations of NC-derived inferior turbinate stem cells from distinct donors. In accordance to our present observations, Clewes and colleagues reported a donor-dependent molecular signature of epidermal neural crest-derived stem cell, particularly including pluripotency-associated genes ^6^. Although the age of the donor is often related to variations in the behavior of adult stem cells (reviewed in ^31^), we could not observe an association between donor age and the heterogeneous protein amounts of Nestin, S100, Slug or NF-κB. In particular, female donors of similar age (female donor II: 67 y, female donor III: 61 y) revealed a significant difference in the amounts of Nestin, S100, Slug and NF-κB protein. In addition, the interpopulational heterogeneity in marker expression shown here was not attributed to potential sexual dimorphisms, which are increasingly noticed to drive heterogeneous behavior of NCSCs and other adult stem cells (^20^ reviewed in ^32^). We therefor suggest genetic variations between the distinct donors to account for the observed differences in NCSC-marker expression between the analyzed ITSC-populations, which will be assessed in more detail in future work.

Next to the donor-dependent variations between distinct ITCSC-populations, we even observed a great intrapopulational heterogeneity of NCSC-marker protein amounts within the analyzed populations. In accordance to our findings, a range of studies reported intrapopulational variations of NCSCs, namely between clonally grown NCSC-cultures from the same population. The NCSC-clones were shown to strongly vary in their differentiation potential and molecular profile (reviewed in ^2^). In particular, Singhatanadgit and coworkers reported a highly heterogeneous differentiation potential between clonal human periodontal ligament stem cell cultures, which varied from sole osteogenic to multilineage differentiation ^33^. Human dental pulp stem cell (DPSC) clones were further reported to reveal a great variability in the potential for generating dentin *in vitro* and *in vivo ^19^*. In previous studies, we also observed clonal ITSC-cultures to reveal different ratios of ectodermal to mesodermal progeny upon spontaneous differentiation ^13,22^. Linking this genetic heterogeneity in differentiation potential of NCSCs to heterogeneous gene expression levels of NCSC-markers, Young and colleagues demonstrated Nestin to be highly differentially expressed between murine DPSC-clones. The authors further observed a high expression of Nestin to be a pre-requisite for neuronal and oligodendrocyte differentiation ^34^. Extending these promising findings, our present results show highly heterogenous protein amounts of Nestin, S100, Slug and NF-κB-p65 even between single ITSCs in the same clonal culture. In addition to intrinsic genetic variations among clonal cultures, these observations suggest stochastic heterogeneity to at least partly account for the variability in NCSC-marker expression shown between single ITSCs. Stochastic variations in transcription or cell cycle have been already shown to account for heterogeneous transcription factor activity, molecular profiles and behavior of embryonic stem cells (ESCs)^35,36,37^. Such stochastic transcriptional heterogeneity of ESCs can be particularly based on histone modifications^38^, allele-switching^36^, mRNA half-life^36^ and transcriptional bursting, the stochastic activation and inactivation of promoters ^39^, which is in turn driven by a range of promoter- and gene body-binding proteins^40^. Prost-transcriptional variations in protein synthesis and degradation further contribute to the regulation of heterogeneous stem cell identities and enable stem cell homeostasis^41^. Among adult stem cells, heterogeneity of hematopoietic stem cells (HSCs) is also controversially discussed to be partly driven by stochastic variations^42–44^. Stochastic fluctuations in transcription as discussed above as well as bistable epigenetic states are also suggested to build the basis of intraclonal heterogeneity observed in mesenchymal stem cells (MSCs)^45^(reviewed in ^31^). Despite the well-studied basis of stochastic heterogeneity in ESCs and their potential role in adult HSCs and MSCs, similar mechanisms remain to be investigated in more detail in human neural crest-derived stem cells. The present study provides evidence for the first time that intrapopulational heterogeneity of NC-derived human ITSCs is at least partly based on stochastic variations in NCSC-markers, which may include variations in cell cycle or transcription as discussed above. We propose this stochastic intrapopulational heterogeneity to occur in addition to the already described genetic variability between clonal NCSC-cultures and the niche-dependent plasticity of NCSCs (reviewed in ^2^). Off note, extrinsic stimuli related to the culture condition may be suggested to likewise account for the heterogeneous expression of NCSC-markers. However, the cultivation method utilized here was previously demonstrated to assure genetic stability and stemness of ITSCs ^22^, thus minimizing potential extrinsic influences.

On regulatory level, our data show an increased nuclear protein amount of Slug in ITSCs from one female donor, which was accompanied by a significantly elevated level of nuclear NF-κB-p65 protein, suggesting an NF-κB-dependent regulation of Slug. In this line, Zhang and coworkers reported Slug to be directly regulated by NF-κB in the early vertebrate mesoderm, while Slug upregulation in turn indirectly drove NF-κB RelA expression^26^. Slug belongs to a group of transcription factors also including amongst others SNAIL and TWIST, which are required for neural crest formation and enable EMT and therefore migration of embryonic NCCs ^26,46^. For adult NCSCs, we recently suggested such NC-related transcription factors as Slug as well as NF-κB to be vital for maintaining stemness (reviewed in ^2^). Accordingly, Tang and coworkers demonstrated SNAIL and Slug to cooperatively control self-renewal and differentiation of adult stem cells from the murine bone marrow47. We further recently reported inhibition of NF-κB c-Rel during differentiation of ITSCs to result in a fate shift from the neuronal to oligodendroglial lineage^30^. The present data extend these promising findings by demonstrating heterogeneous expression and protein amounts of these core NCSC-markers as well as NF-κB on an inter- and intrapopulational level, suggesting the presence of heterogeneous NCSC-stemness states. Accordingly, Gadye and coworkers reported the presence of transient stemness states in murine NCSCs residing in the olfactory epithelium following injury ^48^.

In summary, we show a donor-dependent interpopulational variation in NCSC-marker expression between ITSC-populations from distinct donors. This interpopulational heterogeneity was accompanied by stochastic intrapopulational differences in NCSC-marker proteins between single ITSCs, which we consider as an additional level of heterogeneity to the already described genetic heterogeneity between clonal NCSC-cultures and their niche-dependent plasticity. Our present findings thus offer a broader perspective on NCSC-heterogeneity and may serve to better understand NCSC-behavior and thus improve their clinical applicability.

## Acknowledgements

This work was funded by the Bielefeld University. BW was funded by an internal grant of the Bethel Foundation, Bielefeld, Germany.

## Conflicts of interest/Competing interests

The authors declare no conflict of interest or competing interests.

## Ethics approval

All experimental procedures were ethically approved by the ethics board of the medical faculty of the University of Münster (No. 2012–015-f-S).

## Consent to participate and for publication

Inferior turbinate stem cells were isolated from human inferior turbinate tissue obtained during routine nasal surgery according to Hauser and coworkers^13^ after an informed consent according to local and international guidelines.

## Availability of data and material (data transparency)

All data are made available within the manuscript.

## Code availability (software application or custom code)

Not applicable.

## Authors’ contributions

Conceptualization and study design: JG; Formal analysis: BW, AH, JG; investigation: BW, AH; Writing - original draft preparation: AH, JG; Writing - review and editing: BW; Supervision: JG

## Notes

### Competing Interest Statement

The authors have declared no competing interest.

## References

1 His, W. Untersuchungen über die erste Anlage des Wirbeltierleibes. Die erste Entwicklung des Hühnchens im Ei. (Vogel, 1868).

2 Höving, A. L. et al. Between Fate Choice and Self-Renewal-Heterogeneity of Adult Neural Crest-Derived Stem Cells. Front Cell Dev Biol 9, doi:ARTN 662754 10.3389/fcell.2021.662754 (2021).

3 Kaltschmidt, B., Kaltschmidt, C. & Widera, D. Adult craniofacial stem cells: sources and relation to the neural crest. Stem cell reviews 8, 658–671, doi:10.1007/s12015-011-9340-9 (2012).

4 Dupin, E. & Coelho-Aguiar, J. M. Isolation and differentiation properties of neural crest stem cells. Cytometry A 83, 38–47, doi:10.1002/cyto.a.22098 (2013).

5 Toma, J. G. et al. Isolation of multipotent adult stem cells from the dermis of mammalian skin. Nature cell biology 3, 778–784, doi:10.1038/ncb0901-778 [doi] ncb0901-778 [pii] (2001).

6 Clewes, O. et al. Human epidermal neural crest stem cells (hEPI-NCSC)--characterization and directed differentiation into osteocytes and melanocytes. Stem cell reviews 7, 799–814, doi:10.1007/s12015-011-9255-5 (2011).

7 Pardal, R., Ortega-Saenz, P., Duran, R. & Lopez-Barneo, J. Glia-like stem cells sustain physiologic neurogenesis in the adult mammalian carotid body. Cell 131, 364–377, doi:10.1016/j.cell.2007.07.043 (2007).

8 Höving, A. L. et al. Transcriptome Analysis Reveals High Similarities between Adult Human Cardiac Stem Cells and Neural Crest-Derived Stem Cells. Biology (Basel) 9, doi:10.3390/biology9120435 (2020).

9 Höving, A. L. et al. Blood Serum Stimulates p38-Mediated Proliferation and Changes in Global Gene Expression of Adult Human Cardiac Stem Cells. Cells 9, doi:10.3390/cells9061472 (2020).

10 Widera, D. et al. Highly efficient neural differentiation of human somatic stem cells, isolated by minimally invasive periodontal surgery. Stem Cells Dev 16, 447–460, doi:10.1089/scd.2006.0068 [doi] (2007).

11 Murrell, W. et al. Olfactory mucosa is a potential source for autologous stem cell therapy for Parkinson’s disease. Stem Cells 26, 2183–2192, doi:10.1634/stemcells.2008-0074 (2008).

12 Schurmann, M. et al. Identification of a Novel High Yielding Source of Multipotent Adult Human Neural Crest-Derived Stem Cells. Stem cell reviews, doi:10.1007/s12015-017-9797-2 (2017).

13 Hauser, S. et al. Isolation of novel multipotent neural crest-derived stem cells from adult human inferior turbinate. Stem Cells Dev 21, 742–756 (2012).

14 Tabakow, P. et al. Functional regeneration of supraspinal connections in a patient with transected spinal cord following transplantation of bulbar olfactory ensheathing cells with peripheral nerve bridging. Cell transplantation, doi:10.3727/096368914X685131 (2014).

15 Müller, J. et al. Intrastriatal transplantation of adult human neural crest-derived stem cells improves functional outcome in parkinsonian rats. Stem cells translational medicine 4, 31–43, doi:sctm.2014-0078 [pii] 10.5966/sctm.2014-0078 (2015).

16 Rodrigues, R. M. et al. Human skin-derived stem cells as a novel cell source for in vitro hepatotoxicity screening of pharmaceuticals. Stem Cells Dev 23, 44–55, doi:10.1089/scd.2013.0157 (2014).

17 Müller, J. et al. 1,8-Cineole potentiates IRF3-mediated antiviral response in human stem cells and in an ex vivo model of rhinosinusitis. Clin Sci (Lond) 130, 1339–1352, doi:10.1042/CS20160218 (2016).

18 Annese, V., Navarro-Guerrero, E., Rodriguez-Prieto, I. & Pardal, R. Physiological Plasticity of Neural-Crest-Derived Stem Cells in the Adult Mammalian Carotid Body. Cell Rep 19, 471–478, doi:10.1016/j.celrep.2017.03.065 (2017).

19 Gronthos, S., Mankani, M., Brahim, J., Robey, P. G. & Shi, S. Postnatal human dental pulp stem cells (DPSCs) in vitro and in vivo. Proceedings of the National Academy of Sciences of the United States of America 97, 13625–13630, doi:10.1073/pnas.240309797 (2000).

20 Ruiz-Perera, L. M. et al. NF-kappaB p65 directs sex-specific neuroprotection in human neurons. Scientific reports 8, 16012, doi:10.1038/s41598-018-34394-8 (2018).

21 Greiner, J. F. et al. Natural and synthetic nanopores directing osteogenic differentiation of human stem cells. Nanomedicine. 17, 319–328, doi:10.1016/j.nano.2019.01.018 (2019).

22 Greiner, J. F. et al. Efficient animal-serum free 3D cultivation method for adult human neural crest-derived stem cell therapeutics. Eur Cell Mater 22, 403–419 (2011).

23 Perkins, N. D. Integrating cell-signalling pathways with NF-kappaB and IKK function. Nature reviews. Molecular cell biology 8, 49–62, doi:10.1038/nrm2083 (2007).

24 Kaltschmidt, B. & Kaltschmidt, C. NF-kappaB in the nervous system. Cold Spring Harbor perspectives in biology 1, a001271, doi:10.1101/cshperspect.a001271 (2009).

25 Kaltschmidt, B., Greiner, J. F. W., Kadhim, H. M. & Kaltschmidt, C. Subunit-Specific Role of NF-kappaB in Cancer. Biomedicines 6, doi:10.3390/biomedicines6020044 (2018).

26 Zhang, C., Carl, T. F., Trudeau, E. D., Simmet, T. & Klymkowsky, M. W. An NF-kappaB and slug regulatory loop active in early vertebrate mesoderm. PloS one 1, e106, doi:10.1371/journal.pone.0000106 (2006).

27 Picelli, S. et al. Full-length RNA-seq from single cells using Smart-seq2. Nature Protocols 9, 171, doi:10.1038/nprot.2014.006 https://www.nature.com/articles/nprot.2014.006#supplementary-information (2014).

28 Schindelin, J. et al. Fiji: an open-source platform for biological-image analysis. Nature Methods 9, 676–682, doi:10.1038/nmeth.2019 (2012).

29 Pires, B. R. et al. NF-kappaB Is Involved in the Regulation of EMT Genes in Breast Cancer Cells. PloS one 12, e0169622, doi:10.1371/journal.pone.0169622 (2017).

30 Ruiz-Perera, L. M., Greiner, J. F. W., Kaltschmidt, C. & Kaltschmidt, B. A Matter of Choice: Inhibition of c-Rel Shifts Neuronal to Oligodendroglial Fate in Human Stem Cells. Cells 9, doi:10.3390/cells9041037 (2020).

31 McLeod, C. M. & Mauck, R. L. On the origin and impact of mesenchymal stem cell heterogeneity: new insights and emerging tools for single cell analysis. Eur Cell Mater 34, 217–231, doi:10.22203/eCM.v034a14 (2017).

32 Greiner, J., Merten, M., Kaltschmidt, C. & Kaltschmidt, B. Sexual dimorphisms in adult human neural, mesoderm-derived, and neural crest-derived stem cells. FEBS Lett, doi:10.1002/1873-3468.13606 (2019).

33 Singhatanadgit, W., Donos, N. & Olsen, I. Isolation and characterization of stem cell clones from adult human ligament. Tissue Eng Part A 15, 2625–2636, doi:10.1089/ten.TEA.2008.0442 (2009).

34 Young, F. I. et al. Clonal Heterogeneity in the Neuronal and Glial Differentiation of Dental Pulp Stem/Progenitor Cells. Stem Cells Int 2016, 1290561, doi:10.1155/2016/1290561 (2016).

35 Huh, D. & Paulsson, J. Non-genetic heterogeneity from stochastic partitioning at cell division. Nat Genet 43, 95–100, doi:10.1038/ng.729 (2011).

36 Torres-Padilla, M.-E. & Chambers, I. Transcription factor heterogeneity in pluripotent stem cells: a stochastic advantage. Development 141, 2173–2181, doi:10.1242/dev.102624 (2014).

37 Wu, J. & Tzanakakis, E. S. Contribution of Stochastic Partitioning at Human Embryonic Stem Cell Division to NANOG Heterogeneity. PloS one 7, e50715, doi:10.1371/journal.pone.0050715 (2012).

38 Graf, T. & Stadtfeld, M. Heterogeneity of embryonic and adult stem cells. Cell Stem Cell 3, 480–483, doi:10.1016/j.stem.2008.10.007 (2008).

39 Raj, A., Peskin, C. S., Tranchina, D., Vargas, D. Y. & Tyagi, S. Stochastic mRNA Synthesis in Mammalian Cells. PLOS Biology 4, e309, doi:10.1371/journal.pbio.0040309 (2006).

40 Ochiai, H. et al. Genome-wide kinetic properties of transcriptional bursting in mouse embryonic stem cells. Sci Adv 6, eaaz6699, doi:10.1126/sciadv.aaz6699 (2020).

41 Chua, B. A., Van Der Werf, I., Jamieson, C. & Signer, R. A. J. Post-Transcriptional Regulation of Homeostatic, Stressed, and Malignant Stem Cells. Cell Stem Cell 26, 138–159, doi:10.1016/j.stem.2020.01.005 (2020).

42 Abkowitz, J. L., Catlin, S. N. & Guttorp, P. Evidence that hematopoiesis may be a stochastic process in vivo. Nat Med 2, 190–197, doi:10.1038/nm0296-190 (1996).

43 Muller-Sieburg, C. E., Sieburg, H. B., Bernitz, J. M. & Cattarossi, G. Stem cell heterogeneity: implications for aging and regenerative medicine. Blood 119, 3900–3907, doi:10.1182/blood-2011-12-376749 (2012).

44 Chang, H. H., Hemberg, M., Barahona, M., Ingber, D. E. & Huang, S. Transcriptome-wide noise controls lineage choice in mammalian progenitor cells. Nature 453, 544–547, doi:10.1038/nature06965 (2008).

45 Hamidouche, Z. et al. Bistable Epigenetic States Explain Age-Dependent Decline in Mesenchymal Stem Cell Heterogeneity. Stem Cells 35, 694–704, doi:10.1002/stem.2514 (2017).

46 Casas, E. et al. Snail2 is an essential mediator of Twist1-induced epithelial mesenchymal transition and metastasis. Cancer research 71, 245–254, doi:10.1158/0008-5472.CAN-10-2330 (2011).

47 Tang, Y., Feinberg, T., Keller, E. T., Li, X. Y. & Weiss, S. J. Snail/Slug binding interactions with YAP/TAZ control skeletal stem cell self-renewal and differentiation. Nature cell biology 18, 917–929, doi:10.1038/ncb3394 (2016).

48 Gadye, L. et al. Injury Activates Transient Olfactory Stem Cell States with Diverse Lineage Capacities. Cell Stem Cell 21, 775–790 e779, doi:10.1016/j.stem.2017.10.014 (2017).

